# The Role of Carbon Nanoparticle in Lymph Node Detection and Parathyroid Gland Protection during Thyroidectomy- a Meta Analysis

**DOI:** 10.1101/783993

**Authors:** Shao-Wei Xu, Zhi-Feng Li, Man-Bin Xu, Han-Wei Peng

## Abstract

**Objective:** To assess the efficiency of the carbon nanoparticles (CNs) in lymph node identification and parathyroid gland protection during thyroidectomy.

**Methods:** A systematic literature search for relevant literatures published up to December 2018 in PubMed, Embase, Web of Science and Cochrane Library was performed. Both English and Chinese literatures were retrieved and analyzed. Randomized controlled trials or nonrandomized controlled trials on the use of CNs during thyroidectomy were enrolled in this study. The primary outcomes included the number of lymph nodes harvesed, rate of lymph nodes involvement, and the rates of accidental parathyroidectomy, hypoparathyroidism, and hypocalcemia. Weighted mean differences (WMDs), odds ratios (ORs) and risk differences (RDs) were calculated for the dichotomous outcome variables. Between-study heterogeneity was tested using the *Q* tests and the *I*^*2*^ statistics. All analyses were performed using Review Manager (version 5.3.5).

**Results:** 25 studies comprising 3266 patients were included in this analysis. The total number of lymph nodes harvested in the groups of carbon nanoparticles was significant higher than that in the control groups (WMD, 2.36; 95% CI, 1.40 to 3.32; *P* <0.01). Administrating carbon nanoparticles was associated with a lower incidence of accidental parathyroid gland removal (OR = 0.30, 95% CI = 0.23 to 0.40, *P* <0.01) and lower rates of both postoperative transient hypoparathyroidism (OR =0.46, 95% CI = 0.33 to 0.64, *P* <0.01) and transient hypocalcemia (OR =0.55, 95% CI = 0.09 to 3.43, *P* =0.52). There was no significant difference of identified lymph node metastatic rates between the patients with and without use of carbon nanoparticles. Subgroup analyses indicated that the application of CNs in thyroid cancer reoperation also decreased the rate of transient hypoparathyroidism (OR =0.20, 95% CI = 0.36 to 0.04, *P* =0.01) and the possibility of accidental parathyroid glands removal (OR = 0.19, 95% CI = 0.05 to 0.73, P<0.05).

**Conclusions:** The application of CNs for thyroidectomy results in higher number of lymph node harvested and better parathyroid gland protection during initial surgery and reoperation for thyroid cancer.

## Introduction

Thyroid cancer is one of the most common types of cancer in the world, the incidence of which increased dramatically in recent two decades(1). Cervical lymph node metastases are common in patients with papillary thyroid carcinoma (PTC), the most common subtype of thyroid cancer(1). Total thyroidectomy combined with levelⅡ-Ⅵ neck dissection for clinically N1b cases or central compartmental dissection (CND) for cN0 case is widely advocated as one of the standard treatment protocols for thyroid cancer, whereas lobectomy is only accepted for T1 cases without high risk factors(2–4). However, it’s a challenge for a surgeon to perform a total thyroidectomy plus CND due to the potential risk of postoperative hypocalcemia or hypoparathyroidism, whose incidence ranges from20% to 60% based on the previous reports and even higher for those who undergo a re-operation(5, 6).

Carbon nanoparticles (CNs) have been successfully used for sentinel lymph node detection in breast carcinoma, gastric carcinoma, and etc. (7, 8). The CNs, with a diameter of 150 nm, can easily be captured by macrophages and penetrate the lymphatic capillaries with an endothelial cell gap of 20–50 nm rather than the capillary vessels whose endothelial cell gap is 120–500 nm, and they ultimately accumulate in the sentinel lymph nodes. This is the hypothetic basis that CNs can be used as a tracer to detect the sentinel node. In the past decade, CNs had been successfully attempted as a negative developer to protect parathyroid gland during initial thyroidectomy (9). Although a couple of meta-analyses had been published on evaluating the value of CNs in initial thyroid surgery, the data need to be updated (10, 11). Furthermore, there are still doubts about the efficiency of CNs in re-operation thyroidectomy due to the hypothesis that the lymphatic capillaries may be destroyed during initial surgery(12, 13). Therefore, we performed a meta-analysis with more comprehensive and updated studies to summarize the role of application of CNs in lymph node detection and parathyroid gland protection during both initial and completion thyroidectomy.

## Materials and Methods

### Search strategy

Two authors conducted independently a search for relevant literatures up to December 2018 in PubMed, Embase, Web of Science, and Cochrane Library. The following strategy and keywords were used: (carbon) or (carbon nanoparticle) or (nano-carbon) or (carbon particle) or (lymph node tracer) or (lymphatic tracer) and (thyroid). We manually searched the reference lists of eligible studies and Clinical Trials.gov to ensure identification of relevant published and unpublished studies.

### Inclusion criteria

Studies included in the meta-analysis need to fulfill the following criteria: (1) human thyroid carcinoma confirmed by pathology; (2) patients underwent thyroidectomy/lobectomy and/or neck dissection; (3) studies designed to compare the use of CNs with the use of methylene blue (MB) or with blank control; (4) studies on human beings; (5) all the operation in study should be performed by the same medical team/ surgeon; (6) full text available in English or Chinese.

### Exclusion Criteria

Studies were excluded if they (1) have a sample size less than 15, (2) included pregnancy or adolescent(aged<16), (3) provided incomplete data, or (4) included patients with benign thyroid diseases that can’t be separated from thyroid cancers and analyzed.

### Data Extraction

Two reviewers (SW Xu and ZF Li) independently performed first-stage screening of titles and abstracts based on the research question. For the second screening, we retrieved articles in full text according to the initial screening. And then we reviewed the full texts of included studies and recorded the following data independently: first author, year of publication, sample size, description of study population (age), study design (prospective or retrospective), surgical procedure (CNs injection dose, points and waiting time), lymph nodes (number of harvested and positive), parathyroid function (number of identification, incidentally resection and patients with hypoparathyroidism, hypocalcemia) and so on. Any discrepancies were resolved by discussion or referred to the corresponding author (HW Peng).

### Quality Assessment

Concerning the quality of study design, the RCT was assessed according to the Jadad scoring system, which consists of 3 items: randomization (0-2 points), blinding (0-2 points), and descriptions of the withdrawals and dropouts (0 or 1 point). The total possible score was 5 points. With regard to NRCTs, the Newcastle-Ottawa Scale was used.

### Statistical Analysis

All analyses with their 95% confidence intervals (95% CIs) were performed in the current meta-analysis using RevMan 5.3.5(free software downloaded from http://www.cochrane.org). Weighted mean differences (WMDs) were calculated for the continuous outcome variables, and odds ratios (ORs) and risk differences (RDs) were calculated for the dichotomous outcome variables. *Q* tests and *I*^2^ statistics were used to assess the degree of heterogeneity between studies. *P* value less than 0.1 for the *Q* test and an *I*^2^ higher than 50% indicated the existence of significant heterogeneity. A random effects model was used for all analyses.

## Results

### Study Selection and Description

Figure 1 details the study flowchart of the initial search and the subsequent selection of relevant articles. On initial search, 4488 studies were identified through web retrieval in PubMed (n=1153), Embase (n=2801), Web of Science (n=441), Cochrane (n=88) and Clinical Trial.gov (n=5), respectively. A total of 4202 apparently irrelevant references and 230 duplicates were excluded. In addition, 31 studies were excluded due to the reasons listed in the diagram. Finally, 25 studies fulfilled the inclusion criteria for the meta-analysis(13–37).

**Fig 1.**
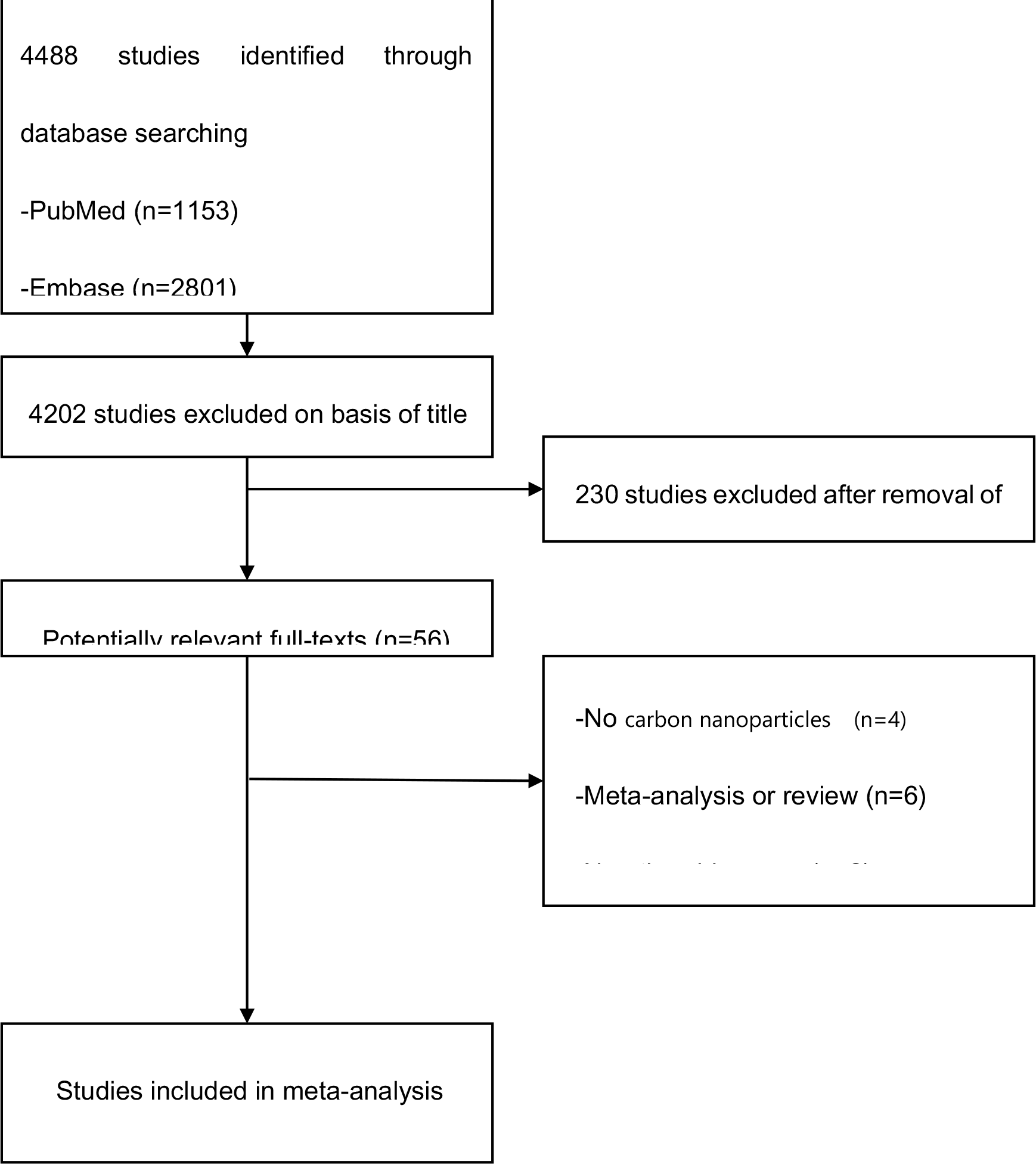
Study flow diagram.

Table 1 summarizes individual studies and their characteristics. A total of 3266 patients were included in this meta-analysis, of which 1496 were included in CN group, 1770 in Control group (1496 in blank control and 181 in MB control), respectively. The two groups had no significant difference in terms of age (MD, −0.43; 95% CI, −1.35-1.26, *P* = 0.95) and sex (OR, 1.05; 95% CI, 0.87-1.25, *P* = 0.63).

**Table 1.**
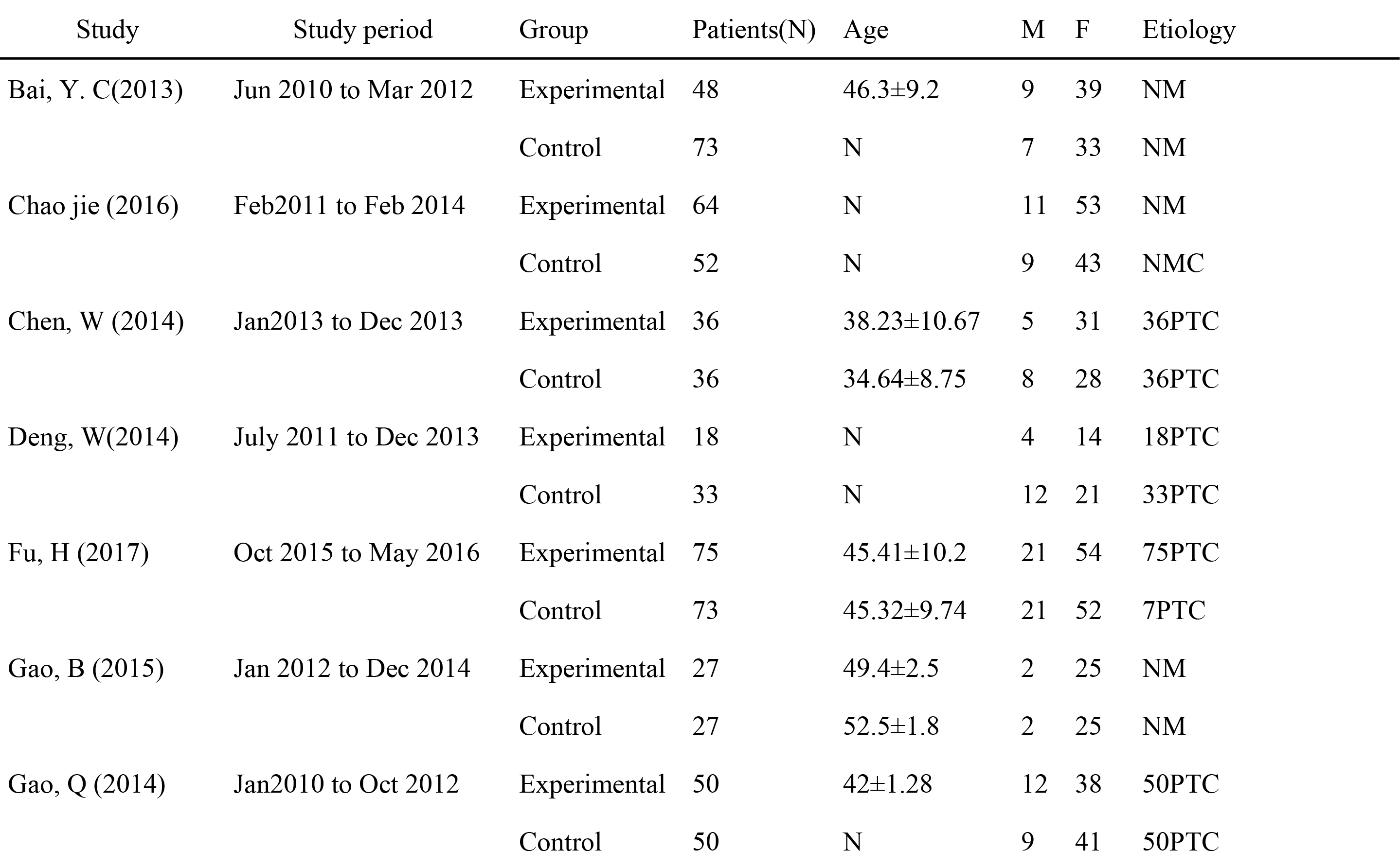
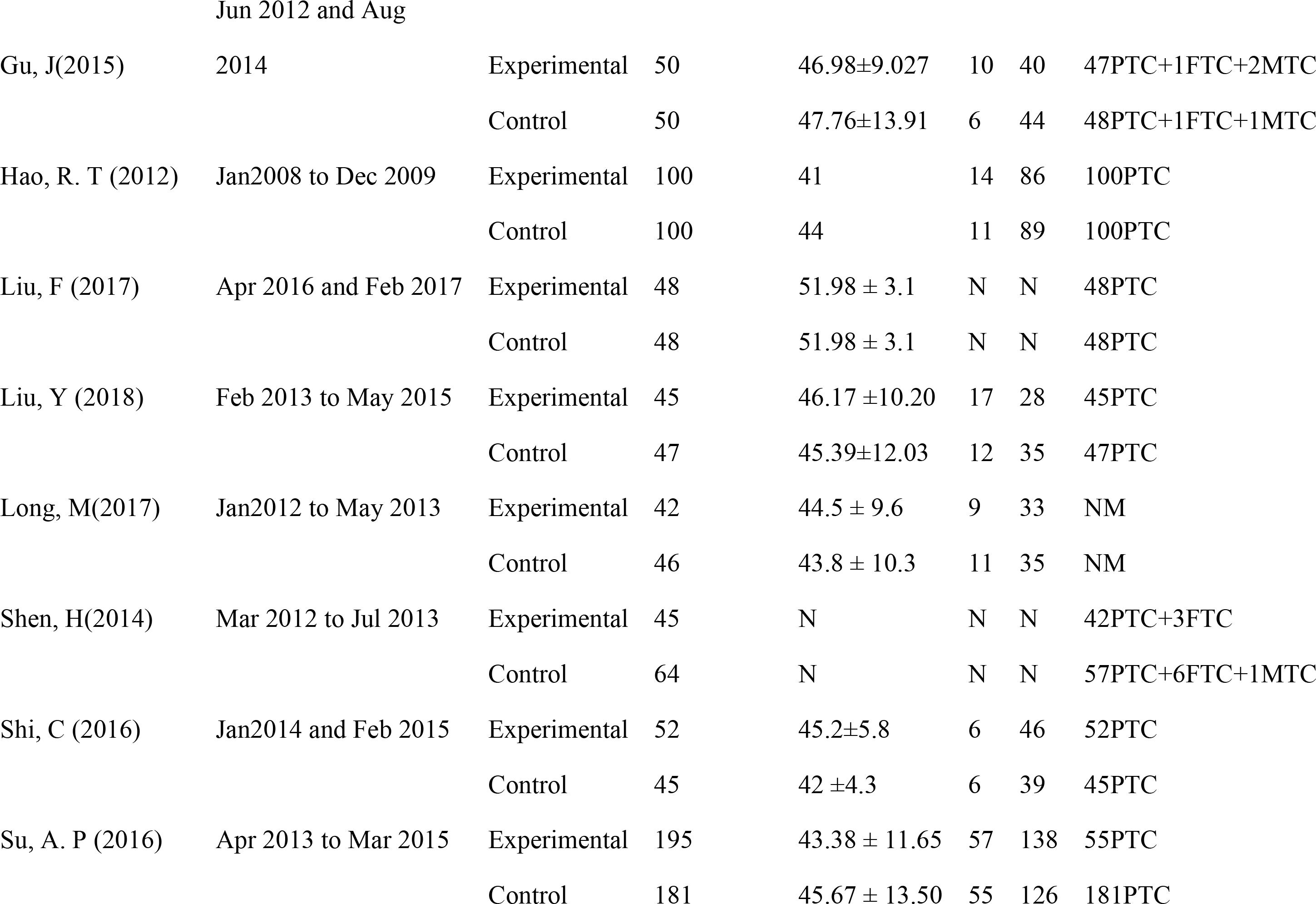
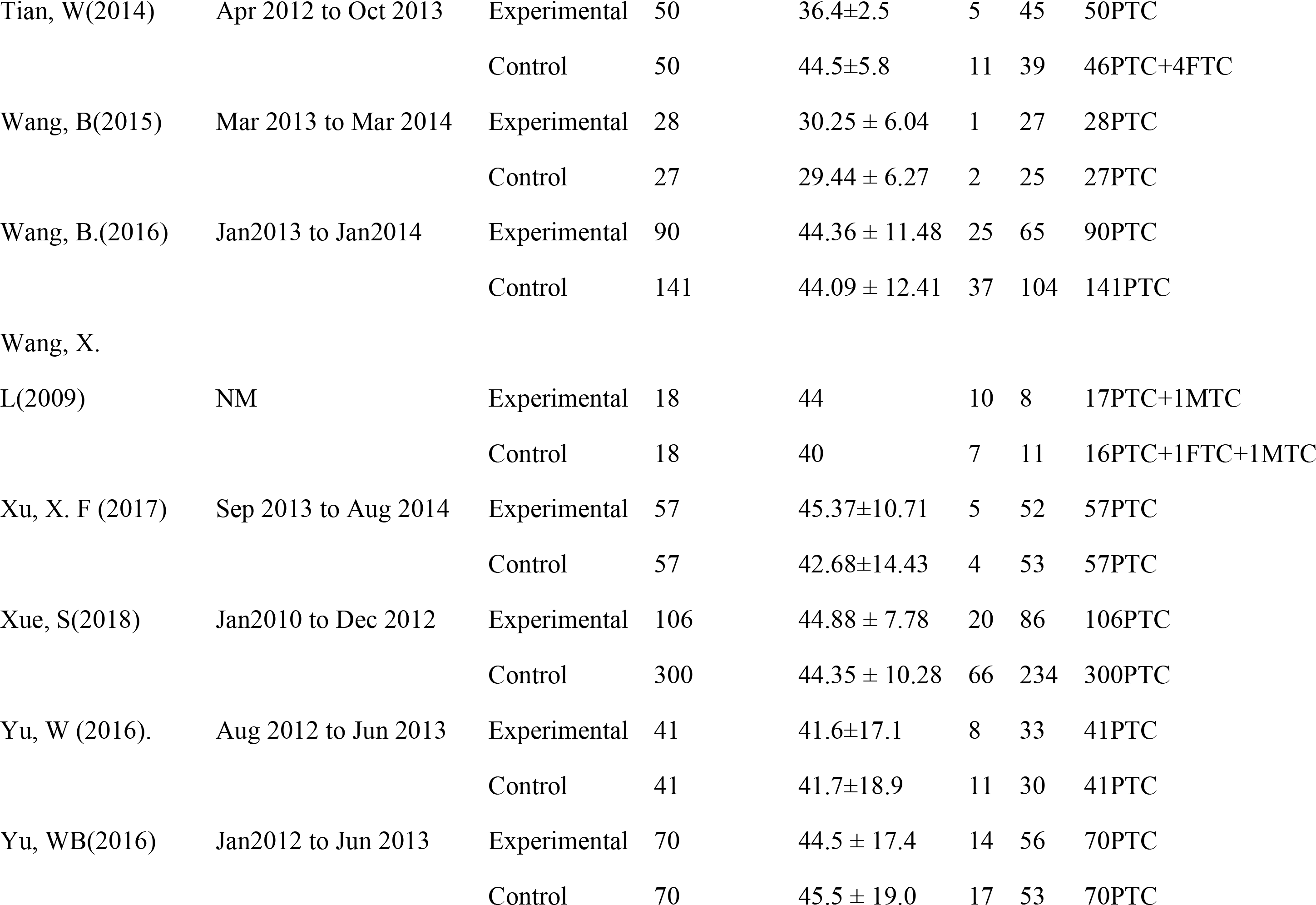
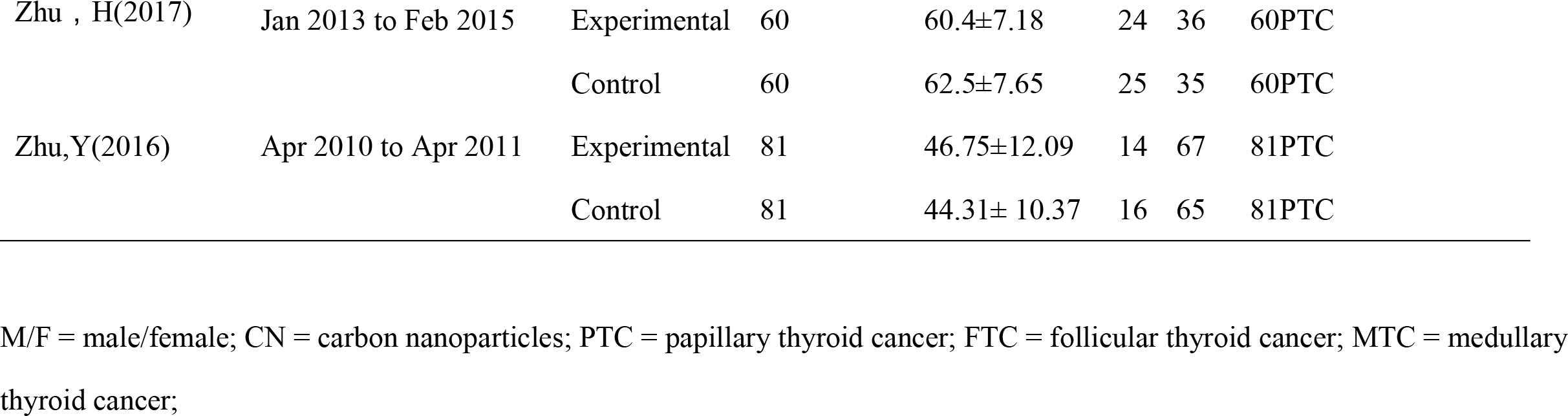
Characteristics of the articles included in the Meta-analysis.

The quality assessment details for the RCTs and NRCTs are presented in Tables 2 and 3.

**Table 2.** Details of Quality Assessment of Randomized Controlled Trials (Jadad scale)

| Study | Randomization | Concealment of Allocation | Blinding | Loss to Follow-up% | Quality Assessment |
| --- | --- | --- | --- | --- | --- |
| Chen, W (2014) | No detailed description | Only mentioned randomized | Unclear | 0 | 2 |
| Gao, B (2015) | No detailed description | Only mentioned randomized | Unclear | 0 | 2 |
| Gao, Q (2014) | No detailed description | Only mentioned randomized | Unclear | 4 | 3 |
| Gu, J(2015) | No detailed description | Only mentioned randomized | Unclear | 0 | 2 |
| Liu, F (2017) | No detailed description | Only mentioned randomized | Unclear | 0 | 2 |
| Long, M(2017) | Computer-generated sequence | Concealed envelopes | Single blinding | 14.6 | 5 |
| Tian, W(2014) | No detailed description | Only mentioned randomized | Unclear | 0 | 2 |
| Wang, B(2015) | Computer-generated<br>sequence | Concealed envelopes | Single blinding | 0 | 5 |
| Xu, X. F (2017) | No detailed description | Only mentioned randomized | Unclear | 0 | 2 |
| Yu, W (2016). | Computer-generated<br>random number tables | Only mentioned randomized | Single blinding | 0 | 4 |
| Yu, WB(2016) | Computer-generated<br>random number tables | Only mentioned randomized | Single blinding | 0 | 4 |
| Zhu, H (2017) | No detailed description | Only mentioned randomized | Unclear | 0 | 2 |
| Zhu, Y(2016) | Computer-generated<br>random number tables | Only mentioned randomized | Single blinding | 0 | 4 |

**Table 3.**
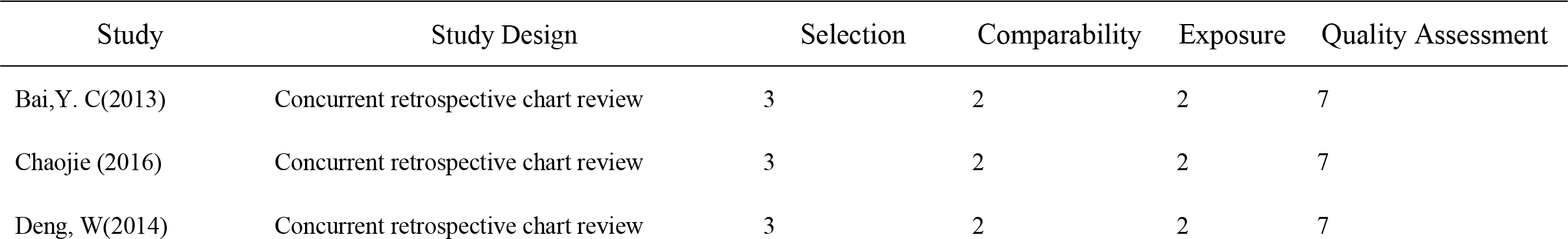
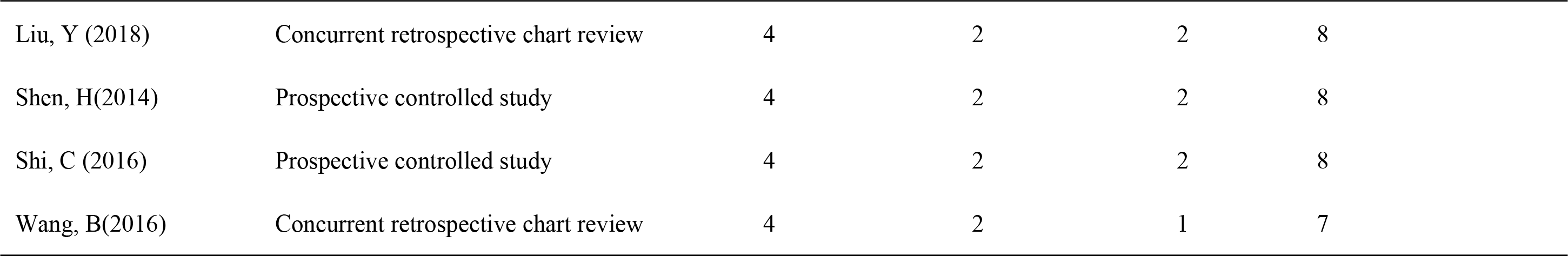
Details of the Quality Assessment of Nocnrandomized Controlled Trial (Newcastle-Ottawa Scale)

### Surgical procedure

The CNs were provided by Chongqing LUMMY Pharmaceutical Co., Ltd. The details of the injection method are shown in Table 4. In most of the studies, the CN injected underneath the fibrous thyroid capsules at two or three points around the neoplasm, 0.1-0.2ml for each point. And 5-10mins later thyroidectomy or lobectomy and neck dissection were performed.

**Table 4.**
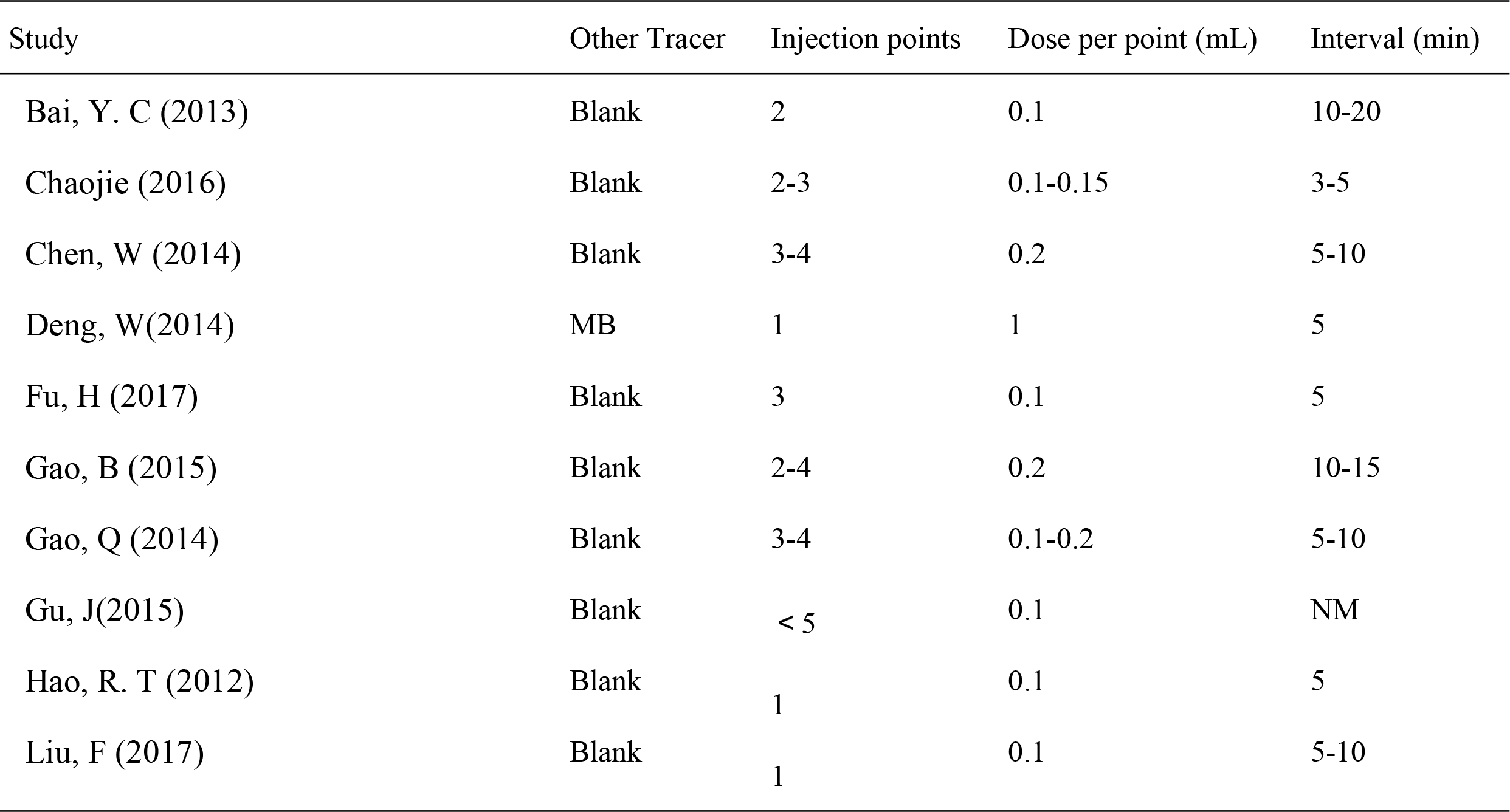
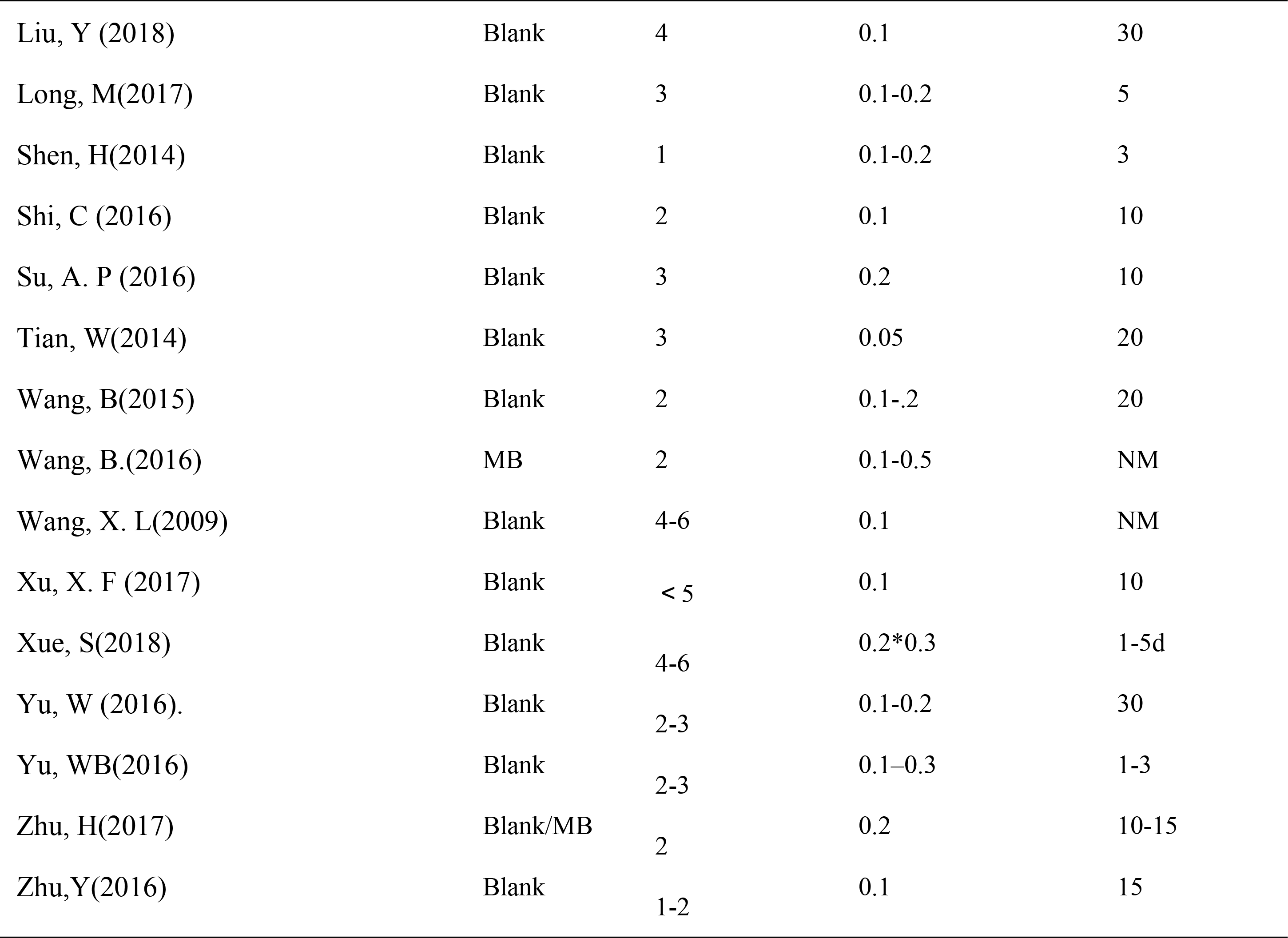
The injection details of carbon nanoparticles used in the Meta-analysis.

### Lymph node removal

All studies regarding lymph node dissection are showed in Figure 2. The total number of lymph node(LN) harvested in the CN group was significant higher than that in the control groups (WMD, 2.36; 95% CI, 1.40 to 3.32; *P*< 0.01, Figure 2). There was no difference in total metastatic rate between the two groups (OR = 1.07, 95% CI = 0.75 to 1.57, *P*=0.71, Figure 3). The rate of LN black-staining varied between 73.3% and 95.3%.

**Fig 2.**
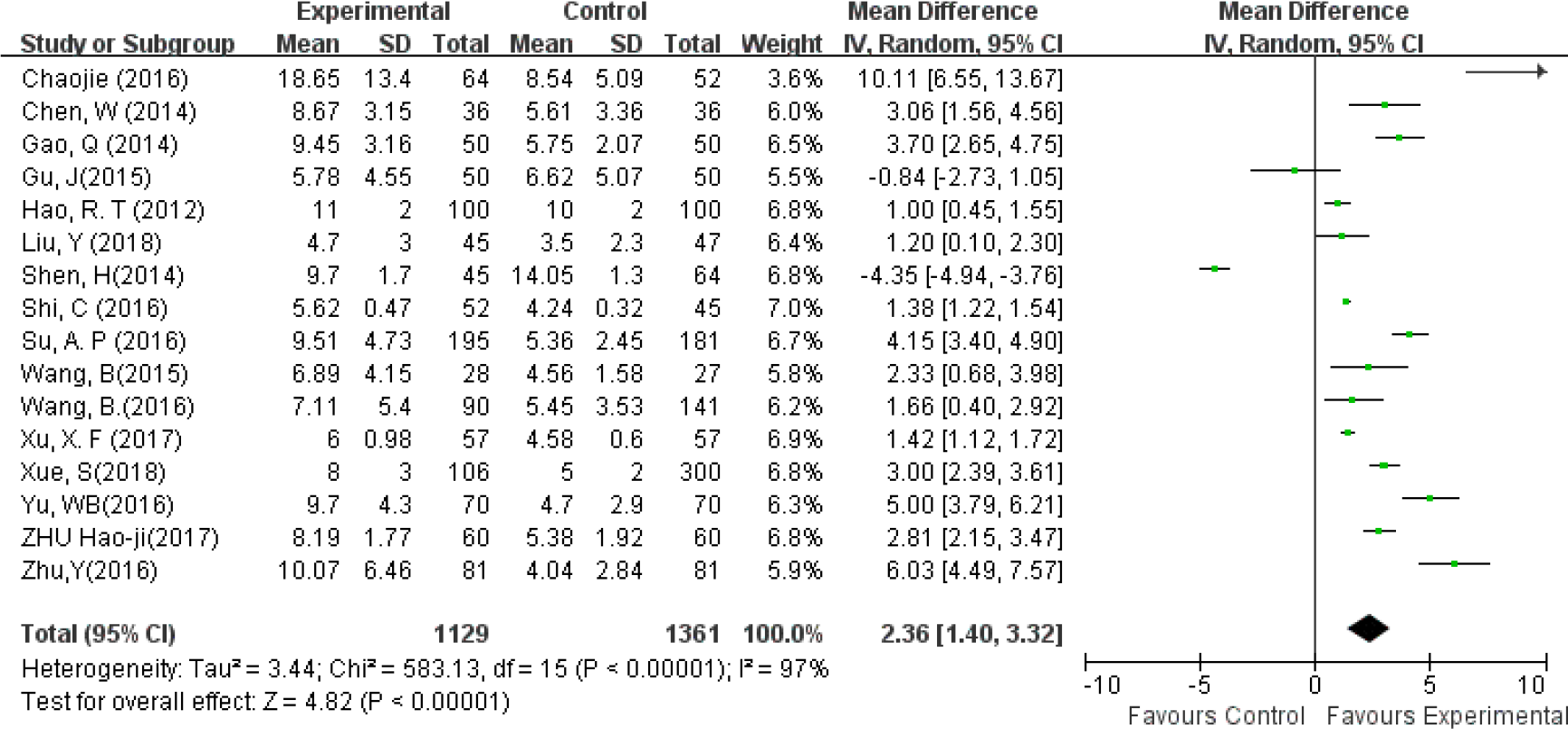
The total number of lymph nodes harvested in groups.

**Fig 3.**
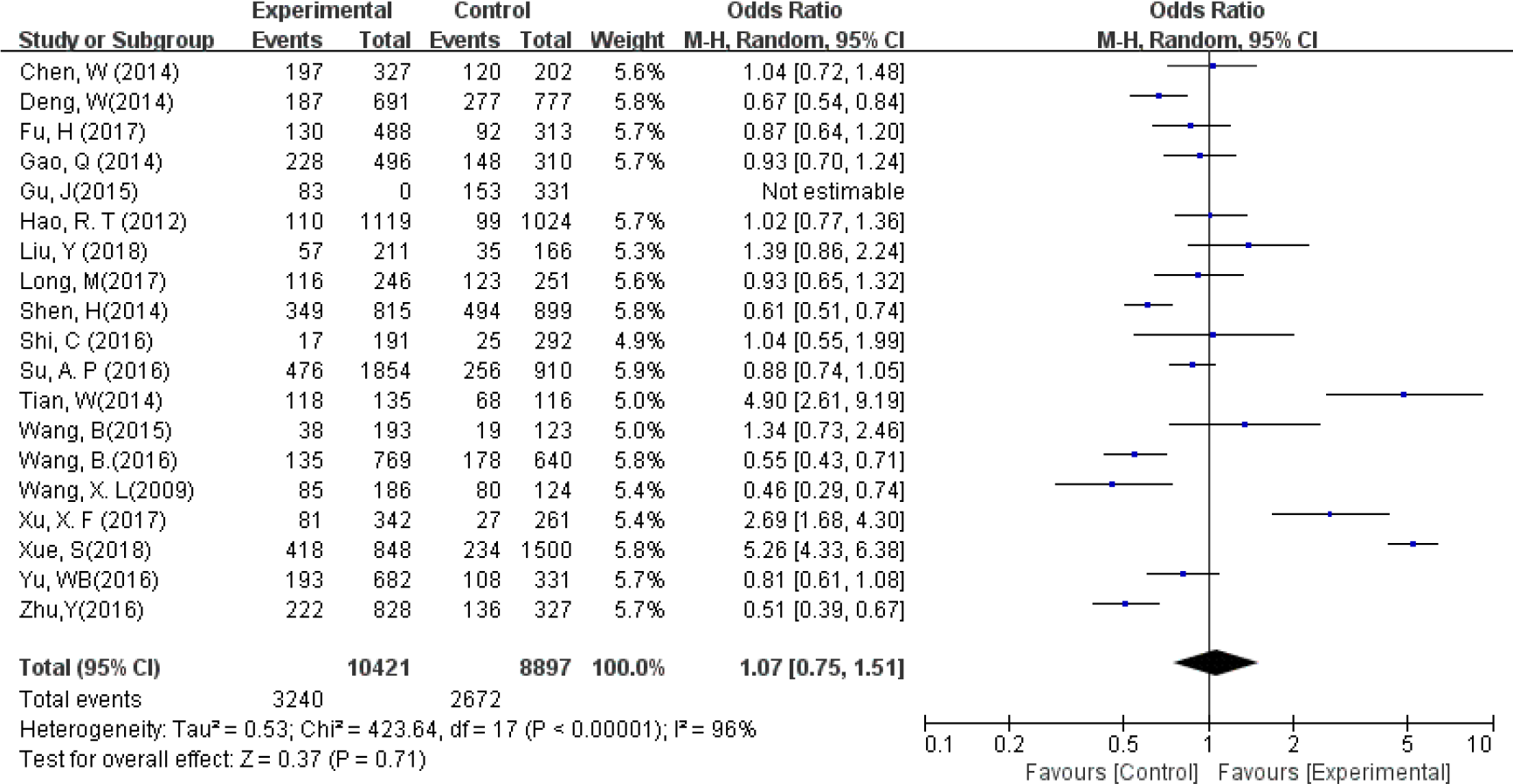
The total metastatic rate of lymph nodes harvested in groups.

### Parathyroid gland protection

The possibility of accidental parathyroid glands (PGs) removal was decreased by 30% in CN group compared with control group (OR = 0.30, 95% CI = 0.23 to 0.40, *P*< 0.01, Figure 4). The application of CN decreased the rate of postoperative transient hypoparathyroidism and transient hypocalcemia by 46% (OR =0.46, 95% CI = 0.33 to 0.64, *P*<0.01, Figure 5a) and 46% (OR =0.46, 95% CI = 0.33 to 0.65, *P* < 0.01,Figure 5b), respectively. The Number of postoperative permanent hypocalcemia in CNs group and control group were 1 and 5, respectively(OR =0.55, 95% CI = 0.09 to 3.43, *P* =0.52,Figure 5c).

**Fig 4.**
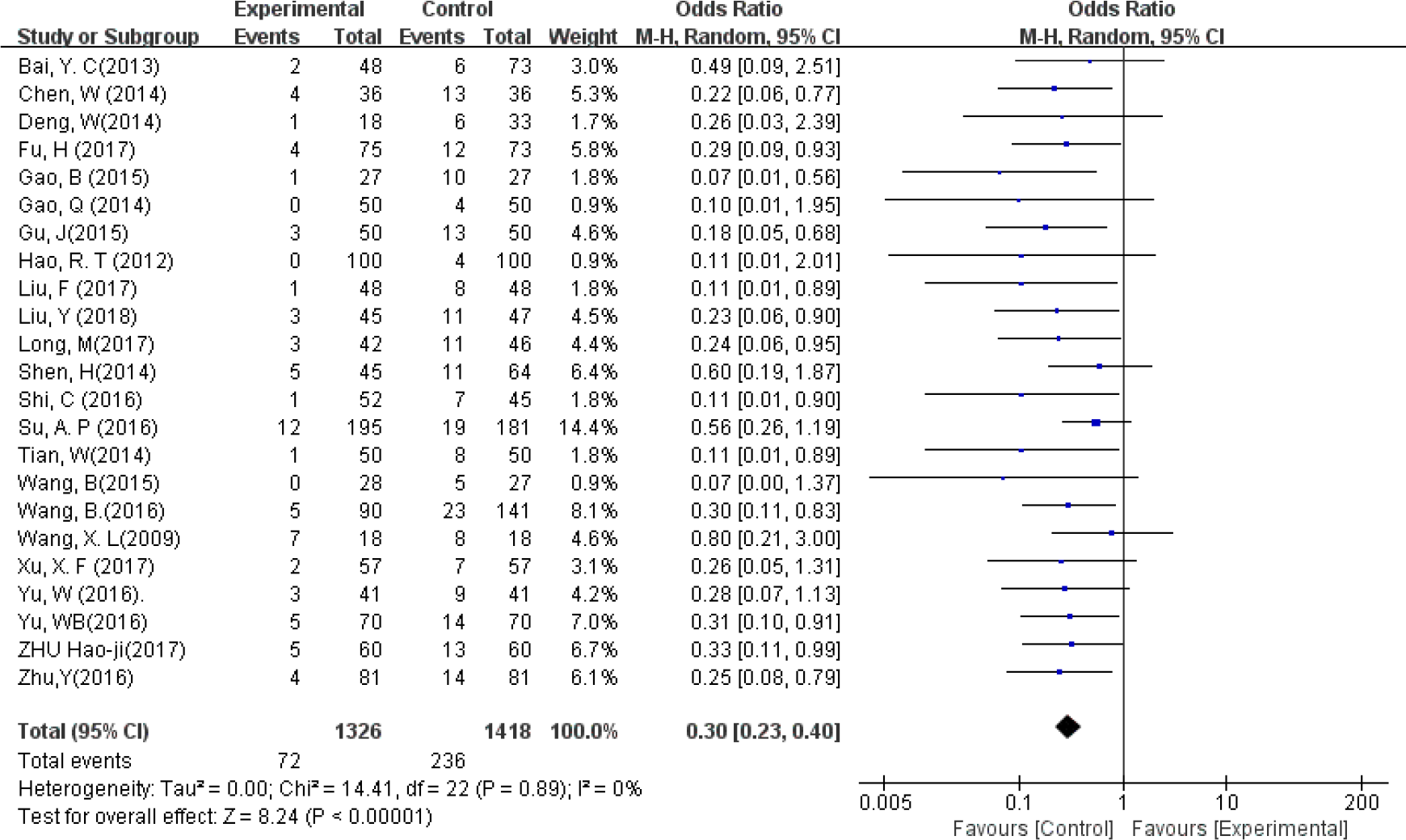
Accidental parathyroid removal rate in groups.

**Fig 5-a.**
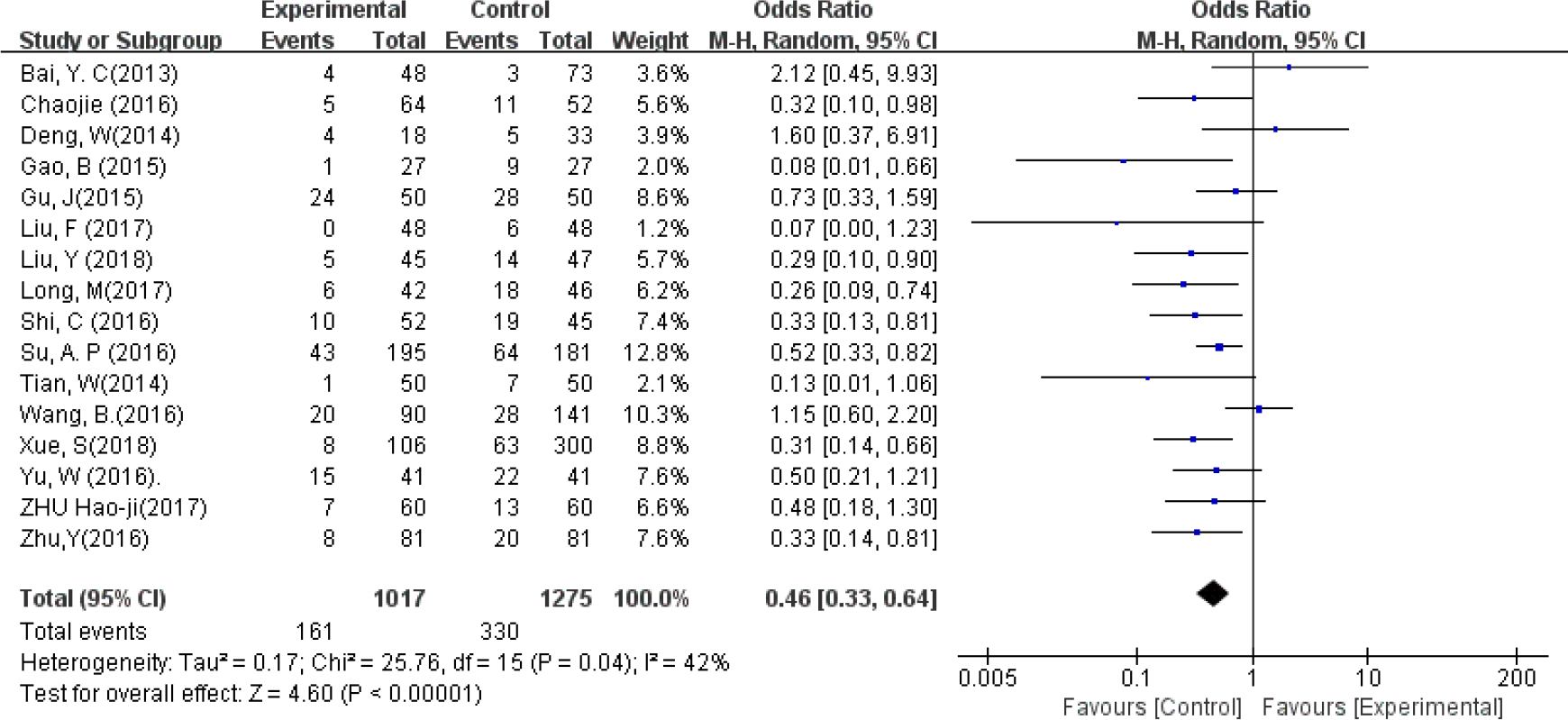
Postoperative transient hypoparathyroidism rate.

**Fig 5-b.**
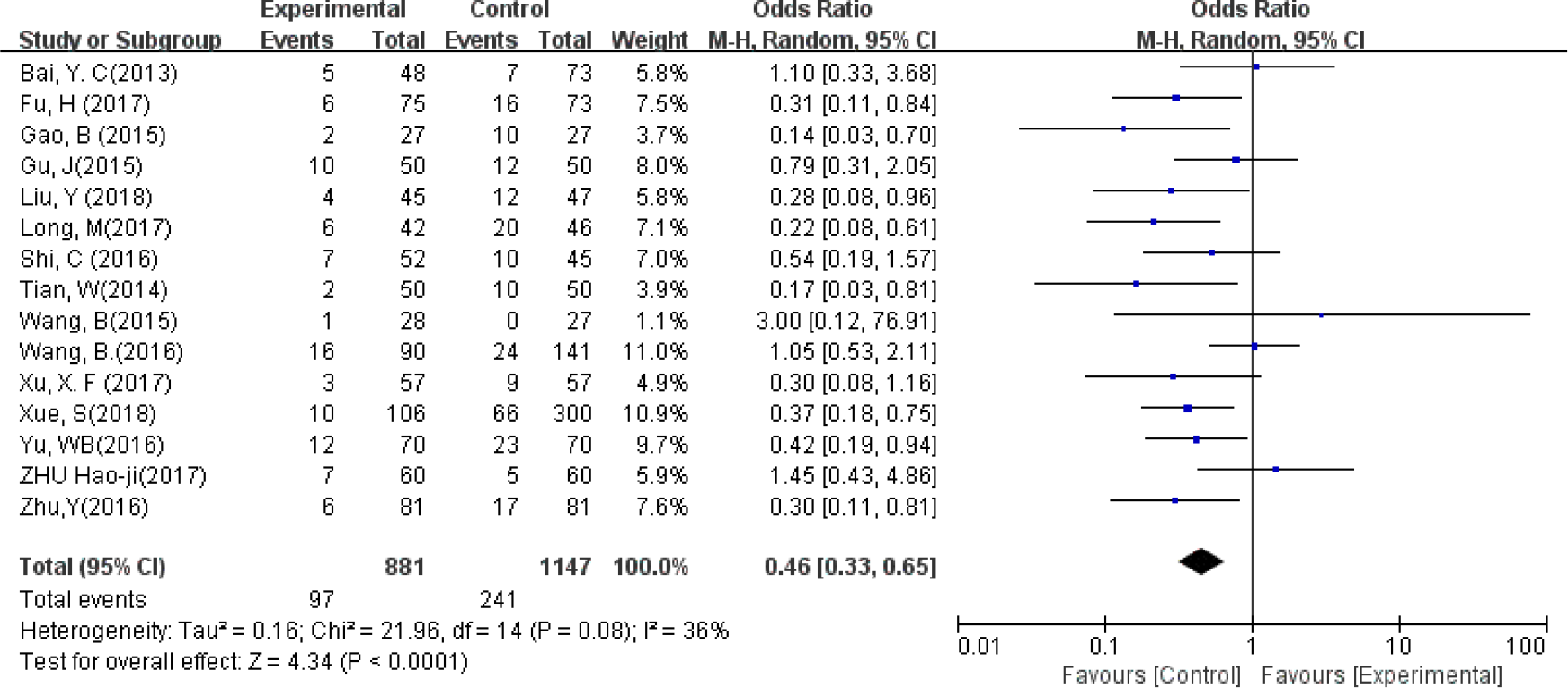
Postoperative transient hypocalcemia rate.

**Fig 5-c.**
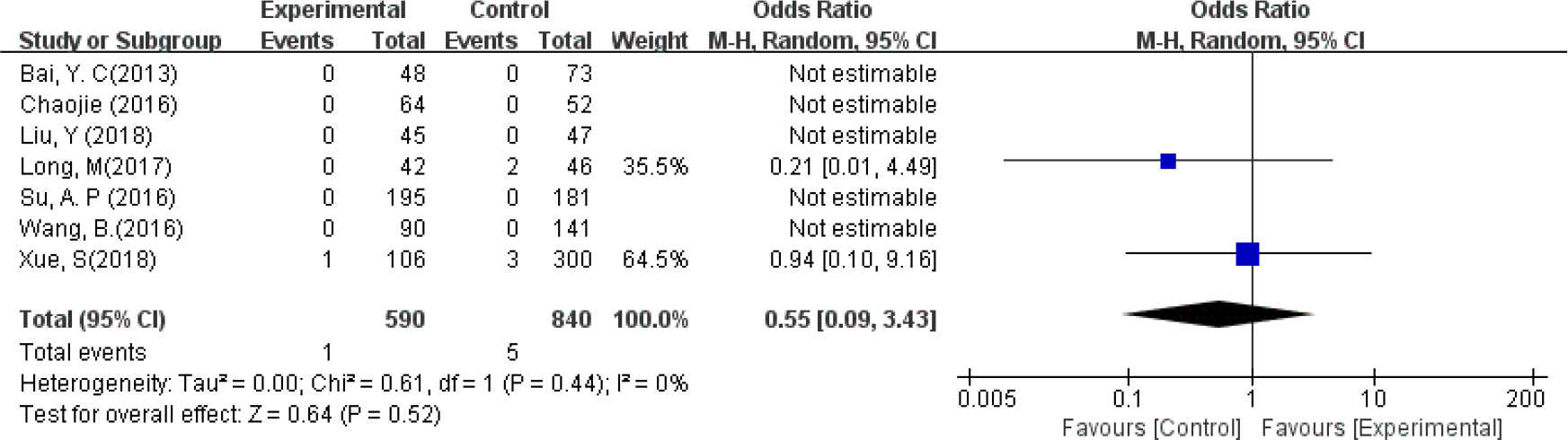
Postoperative permanent hypoparathyroidism rate.

### CNs application during reoperation

For the subgroup of reoperation, CNs significantly decreased the rate of postoperative transient hypoparathyroidism by 21% (OR =0.20, 95% CI = 0.04 to 0.36, *P* =0.01, Figure 6). The possibility of accidental PGs removal was decreased by 19% in CNs group compared with control group during reoperation (OR = 0.19, 95% CI = 0.05 to 0.73, P<0.05, Figure 7).

**Fig 6.**
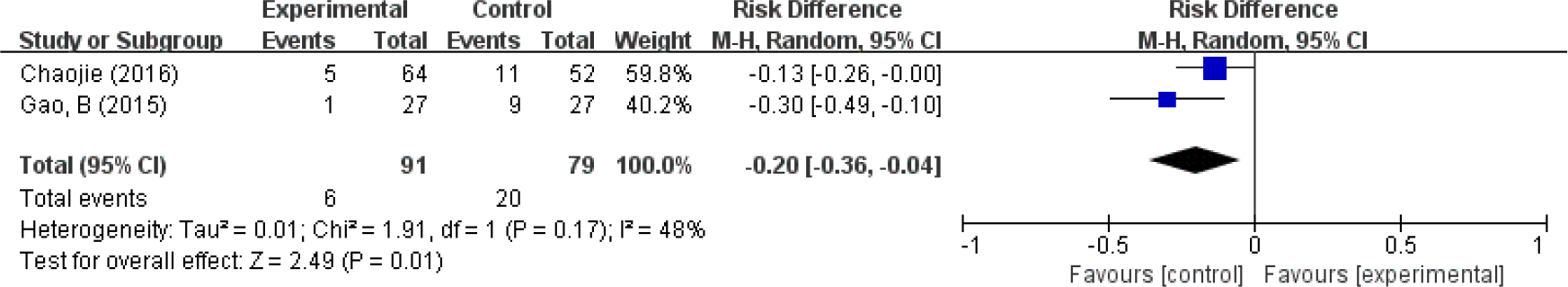
postoperative transient hypoparathyroidism during reoperation.

**Fig 7.**
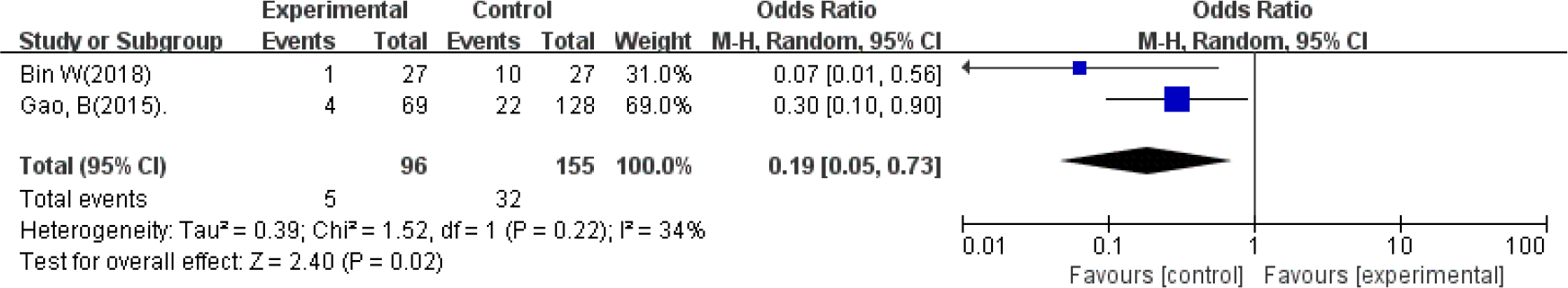
Accidental parathyroid removal rate in groups during reoperation.

## Discussion

Thyroid cancer is one of the most common types of cancer in the world, with a rapid increasing incidence in recent two decades(1). Papillary thyroid carcinoma (PTC), constituting nearly 90% of thyroid cancers, has been reported to have cervical lymph node metastasis in 20%-90% of the patients on their initial diagnosis(1, 38, 39). The most favorite site of metastasis is the central compartment of the neck. CND is recommended for patients with PTC according to most of the current guidelines on the management of thyroid cancers(3, 4). Anatomically, the location of parathyroid glands (PGs), particularly the lower two, are variable, which results in high risk of accidentally removal during thyroidectomy, particularly when a CND is combined(40). Removal of PGs often leads to transient or permanent hypoparathyroidism and hypocalcemia, which has an adverse impact on the quality of life of the patients. Thyroid cancer has a trend of involving youngers and the 5-year survival rate is reported up to 98% (1). As a result, omission of micro-metastases of the central compartmental lymph nodes may lead to reoperation, which may increase incidence of complications and mortality. Therefore, effective techniques are urgently needed to improve the thoroughness of CND while preserving the PGs, especially for reoperation.

Carbon nanoparticles (CN), approved by China Food and Drug Administration, have been well applied to assist lymph node dissection for gastric cancer, breast cancer and etc(7, 8). In the recent years, CN is used as a novel surgical technology for tracer of lymph nodes and localization of parathyroid during thyroid cancer surgery(21). So far, No toxic side effects have been reported in humans. However, when the CNs was injected improperly and they spread out of the thyroid capsule, it may stain the surgical field, which makes surgical procedure more difficult(21). In most of the included studies, the dose of CNs is 0.1 to 0.2 ml per point and two to three points around tumor within the thyroid capsule were recommended. After properly injection, the CN enter the lymphatic capillaries and stain lymph nodes in 10-15min, rather than PGs and the recurrent laryngeal nerve(13–16, 20, 26, 32–37).

Based on this analysis, the total number of lymph node harvested in the group of CN was approximately 2.36 more than that in the control groups. But the ratio of metastatic lymph nodes has no significant differences. The results were consistent with the previous studies. CNs play a key role in accurately identifying lymph nodes, but could not improve the detection of metastatic lymph nodes. As reported that tumor inflammation or injury could block lymphatic channels, lymphatic capillaries around the metastatic lymph nodes may be blocked by tumor cells, resulting in misidentifications because the CN flow is impaired. Application of CN brings convenience to complete lymph nodes detection during neck dissection, but not help distinguish the metastatic lymph nodes.

Hypoparathyroidism is a common complication after thyroid cancer surgery, especially reoperation. The incidence of transient hypoparathyroidism after surgery is reported to be 20% to 60%, while that of permanent hypoparathyroidism is 0% to 7%(5, 6). Our study revealed that application of CNs during lobectomy or CND reduce the rate of PGs accidental removal by approximately 30%. But there were still doubts about practicability for parathyroid protection during thyroid surgery. Liu and Shen reported that CNs play important role in identifying and preserving PGs, but not beneficial for protecting the function of parathyroid gland in thyroid surgery(12, 13). Further analysis found that both rate of postoperative transient hypothyroidism and transient hypocalcemia significantly decreased by 46%. To our knowledge, CNs cannot easily enter the lymphatic vessels for the reoperation, which were destroyed during the initial operation. So the application of CNs during reoperation was never reported in the previous meta-analysis. However, subgroup analysis showed that 2 studies including CNs application during thyroid carcinoma reoperation could not only effectively aid in identifying PGs but also significantly lower incidence of postoperative transient hypoparathyroidism by 20%(25, 35). The application of CNs could remarkably reduce the incidence of the postoperative complication of PGs during thyroid carcinoma operation and reoperation.

The current meta-analysis has some limitations and the results should be interpreted with caution. First, all of studies including in this meta-analysis were performed only in China. No reports for other ethnicities and no publications written in other languages were available. Second, the follow-up period is too short to find out the significant difference in permanent hypocalcemia. Third, the differences between the studies have led to heterogeneity. Further, more standardized randomized controlled trials are required.

## Conclusions

This is the meta-analysis so far, firstly focused on the application of CNs during thyroid carcinoma operation and reoperation. Our results confirmed that the application of CN for thyroidectomy improve the lymph node detection and parathyroid gland protection not only for initial surgery but also reoperation.

## Author Contributions

Shao-Wei Xu, study design, data collection/interpretation, drafting/revision, presentation;

Zhi-Feng Li, study design, data collection/interpretation, drafting/revision;

Man-Bin Xu, data interpretation, drafting/revision, critical review of article;

Han-Wei Peng, authors for correspondence, study design, data interpretation, drafting/revision, critical review of article, final approval.

## Disclosures

### Sponsorships

Youth funds of caner hospital, Shantou University medical college, Guangdong, China. (2018A004).

## References

1. Siegel RL, Miller KD, Jemal A. Cancer statistics, 2019. CA Cancer J Clin. 2019;69(1):7–34.

2. Haugen BR, Alexander EK, Bible KC, Doherty GM, Mandel SJ, Nikiforov YE, et al. 2015 American Thyroid Association Management Guidelines for Adult Patients with Thyroid Nodules and Differentiated Thyroid Cancer: The American Thyroid Association Guidelines Task Force on Thyroid Nodules and Differentiated Thyroid Cancer. Thyroid : official journal of the American Thyroid Association. 2016;26(1):1–133.

3. Haugen BR. 2015 American Thyroid Association Management Guidelines for Adult Patients with Thyroid Nodules and Differentiated Thyroid Cancer: What is new and what has changed? Cancer. 2017;123(3):372–81.

4. Ya M. [Interpretation of the management guidelines for patients with thyroid nodules and differentiated thyroid cancer (2012 Chinese edition)]. Lin chuang er bi yan hou tou jing wai ke za zhi = Journal of clinical otorhinolaryngology, head, and neck surgery. 2013;27(16):917–20.

5. Dedivitis RA, Aires FT, Cernea CR. Hypoparathyroidism after thyroidectomy: prevention, assessment and management. Curr Opin Otolaryngol Head Neck Surg. 2017;25(2):142–6.

6. Shoback DM, Bilezikian JP, Costa AG, Dempster D, Dralle H, Khan AA, et al. Presentation of Hypoparathyroidism: Etiologies and Clinical Features. J Clin Endocrinol Metab. 2016;101(6):2300–12.

7. Sawai K, Hagiwara A, Shimotsuma M, Sakakibara T, Imanishi T, Takemoto Y, et al. Rationale of lymph node dissection for breast cancer--from the viewpoint of analysis of axillary lymphatic flow using activated carbon particle CH40. Gan To Kagaku Ryoho. 1996;23 Suppl 1:30–5.

8. Okamoto K, Sawai K, Minato H, Yada H, Shirasu M, Sakakura C, et al. Number and anatomical extent of lymph node metastases in gastric cancer: analysis using intra-lymph node injection of activated carbon particles (CH40). Jpn J Clin Oncol. 1999;29(2):74–7.

9. Li J, Li X, Wang Z. Negative developing of parathyroid using carbon nanoparticles during thyroid surgery. Gland surgery. 2013;2(2):100–1.

10. Li Y, Jian WH, Guo ZM, Li QL, Lin SJ, Huang HY. A Meta-analysis of Carbon Nanoparticles for Identifying Lymph Nodes and Protecting Parathyroid Glands during Surgery. Otolaryngology--head and neck surgery : official journal of American Academy of Otolaryngology-Head and Neck Surgery. 2015;152(6):1007–16.

11. Su AP, Wei T, Gong YP, Gong RX, Li ZH, Zhu JQ. Carbon nanoparticles improve lymph node dissection and parathyroid gland protection during thyroidectomy: a systematic review and meta-analysis. International journal of clinical and experimental medicine. 2018;11(2):463–73.

12. Liu X, Chang S, Jiang X, Huang P, Yuan Z. Identifying Parathyroid Glands with Carbon Nanoparticle Suspension Does Not Help Protect Parathyroid Function in Thyroid Surgery. Surgical innovation. 2016;23(4):381–9.

13. Shen H, Wei B, Feng S, Zhou Q. [Efficiency of carbon nanoparticles in level VI lymphadenectomy for thyroid carcinoma and prevention of postoperative hypoparathyroidism]. Zhonghua er bi yan hou tou jing wai ke za zhi = Chinese journal of otorhinolaryngology head and neck surgery. 2014;49(10):817–20.

14. Su AP, Wei T, Liu F, Gong YP, Li ZH, Zhu JQ. Use of carbon nanoparticles to improve the dissection of lymph nodes and the identification of parathyroid glands during thyroidectomy for papillary thyroid cancer. International journal of clinical and experimental medicine. 2016;9(10):19529–36.

15. Bai YC, Cheng RC, Hong WJ, Ma YH, Qian J, Zhang JM. [Thyroid lymphography:a new clinical approach for protecting parathyroid in surgery]. Zhonghua er bi yan hou tou jing wai ke za zhi = Chinese journal of otorhinolaryngology head and neck surgery. 2013;48(9):721–5.

16. Xue S, Ren P, Wang P, Chen G. Short and Long-Term Potential Role of Carbon Nanoparticles in Total Thyroidectomy with Central Lymph Node Dissection. Scientific reports. 2018;8(1):11936.

17. Hao RT, Chen J, Zhao LH, Liu C, Wang OC, Huang GL, et al. Sentinel lymph node biopsy using carbon nanoparticles for Chinese patients with papillary thyroid microcarcinoma. Eur J Surg Oncol. 2012;38(8):718–24.

18. Wang B, Qiu NC, Zhang W, Shan CX, Jiang ZG, Liu S, et al. The role of carbon nanoparticles in identifying lymph nodes and preserving parathyroid in total endoscopic surgery of thyroid carcinoma. Surgical endoscopy. 2015;29(10):2914–20.

19. Liu F, Zhu Y, Qian Y, Zhang J, Zhang Y, Zhang Y. Recognition of sentinel lymph nodes in patients with papillary thyroid cancer by nano-carbon and methylene blue. Pakistan journal of medical sciences. 2017;33(6):1485–9.

20. Zhu H-jCM-fZY-f. Protective effect of nano-carbon tracers on the parathyroid glands. Chinese journal of tissue engineering research. 2016;20(30):4476–82.

21. Yu W, Zhu L, Xu G, Song Y, Li G, Zhang N. Potential role of carbon nanoparticles in protection of parathyroid glands in patients with papillary thyroid cancer. Medicine. 2016;95(42):e5002.

22. Yu W, Cao X, Xu G, Song Y, Li G, Zheng H, et al. Potential role for carbon nanoparticles to guide central neck dissection in patients with papillary thyroid cancer. Surgery. 2016;160(3):755–61.

23. Gu J, Wang J, Nie X, Wang W, Shang J. Potential role for carbon nanoparticles identification and preservation in situ of parathyroid glands during total thyroidectomy and central compartment node dissection. International journal of clinical and experimental medicine. 2015;8(6):9640–8.

24. Wang XL, Wu YH, Xu ZG, Ni S, Liu J. Parathyroid glands are differentiated from lymph node by activated-carbon particles. Zhonghua er bi yan hou tou jing wai ke za zhi [Chinese journal of otorhinolaryngology head and neck surgery]. 2009;44(2):136–40.

25. Chaojie Z, Shanshan L, Zhigong Z, Jie H, Shuwen X, Peizhi F, et al. Evaluation of the clinical value of carbon nanoparticles as lymph node tracer in differentiated thyroid carcinoma requiring reoperation. International journal of clinical oncology. 2016;21(1):68–74.

26. Shi C, Tian B, Li S, Shi T, Qin H, Liu S. Enhanced identification and functional protective role of carbon nanoparticles on parathyroid in thyroid cancer surgery: A retrospective Chinese population study. Medicine. 2016;95(46):e5148.

27. Gao Q, Zhao D. [Clinical application of carbon nanoparticles labeled lymph node in cervical lymph node dissection with papillary thyroid cancer staged preoperatively as N0]. Lin chuang er bi yan hou tou jing wai ke za zhi = Journal of clinical otorhinolaryngology, head, and neck surgery. 2014;28(24):1938–40.

28. Deng W, Li H, Chen Y, Gao Y, Huang H, Lin S, et al. [Clinical application of carbon nanoparticles in surgery for papillary thyroid carcinoma in young patients]. Zhonghua er bi yan hou tou jing wai ke za zhi = Chinese journal of otorhinolaryngology head and neck surgery. 2014;49(10):812–6.

29. Zhu Y, Chen X, Zhang H, Chen L, Zhou S, Wu K, et al. Carbon nanoparticle-guided central lymph node dissection in clinically node-negative patients with papillary thyroid carcinoma. Head & neck. 2016;38(6):840–5.

30. Long M, Luo D, Diao F, Huang M, Huang K, Peng X, et al. A Carbon Nanoparticle Lymphatic Tracer Protected Parathyroid Glands During Radical Thyroidectomy for Papillary Thyroid Non-Microcarcinoma. Surgical innovation. 2017;24(1):29–34.

31. Liu Y, Li L, Yu J, Fan YX, Lu XB. Carbon nanoparticle lymph node tracer improves the outcomes of surgical treatment in papillary thyroid cancer. Cancer biomarkers : section A of Disease markers. 2018;23(2):227–33.

32. Tian W, Jiang Y, Gao B, Zhang X, Zhang S, Zhao J, et al. Application of nano-carbon in lymph node dissection for thyroid cancer and protection of parathyroid glands. Medical science monitor : international medical journal of experimental and clinical research. 2014;20:1925–30.

33. Chen W, Lv Y, Xie R, Xu D, Yu J. Application of lymphatic mapping to recognize and protect parathyroid in thyroid carcinoma surgery by using carbon nanoparticles. Lin chuang er bi yan hou tou jing wai ke za zhi [Journal of clinical otorhinolaryngology, head, and neck surgery]. 2014;28(24):1918–20, 24.

34. Xu XF, Gu J. The application of carbon nanoparticles in the lymph node biopsy of cN0 papillary thyroid carcinoma: a randomized controlled clinical trial. Asian journal of surgery / Asian Surgical Association. 2017;40(5):345–9.

35. Gao B, Tian W, Jiang Y, Zhang S, Guo L, Zhao J, et al. Application of carbon nanoparticles for parathyroid protection in reoperation of thyroid diseases. International journal of clinical and experimental medicine. 2015;8(12):22254–61.

36. Wang B, Du ZP, Qiu NC, Liu ME, Liu S, Jiang DZ, et al. Application of carbon nanoparticles accelerates the rapid recovery of parathyroid function during thyroid carcinoma surgery with central lymph node dissection: A retrospective cohort study. International journal of surgery (London, England). 2016;36(Pt A):164–9.

37. Fu H, Zhang ZL, Tang ZN, Liu QL. [Application of carbon nanoparticle in neck lymph node dissection for papillary thyroid carcinoma]. Lin chuang er bi yan hou tou jing wai ke za zhi = Journal of clinical otorhinolaryngology, head, and neck surgery. 2017;31(14):1089–92.

38. Wang J, Liu J, Pan H, Jiang C, Liu S, Zhu Z, et al. Young age increases the risk of lymph node positivity in papillary thyroid cancer patients: a SEER data-based study. Cancer management and research. 2018;10:3867–73.

39. Shaha AR, Zaydfudim V, Noguchi S, Thompson GB, Mitchell BK. The impact of lymph node involvement on survival in patients with papillary and follicular thyroid carcinoma DISCUSSION. Surgery. 2008;144(6):1077–8.

40. Sadowski SM, Fortuny JV, Triponez F. A reappraisal of vascular anatomy of the parathyroid gland based on fluorescence techniques. Gland surgery. 2017;6:S30–S7.

